# OH 89: A newly described ∼1.8-million-year-old hominid clavicle from Olduvai Gorge

**DOI:** 10.1101/2023.02.02.526656

**Authors:** Catherine E Taylor, Fidelis Masao, Jackson K Njau, Agustino Venance Songita, Leslea J Hlusko

**Affiliations:** Human Evolution Research Center, University of California Berkeley – Berkeley, CA, USA; University of Dar es Salaam – Dar es Salaam, Tanzania; Conservation Olduvai Project, – Dar es Salaam, Tanzania; Department of Earth and Atmospheric Sciences, Indiana University – Bloomington, Indiana, USA; The Stone Age Institute – Bloomington, Indiana, USA; Centro Nacional de Investigación sobre la Evolución Humana – Burgos, Spain

## Abstract

**Objectives:** Here, we describe the morphology and geologic context of OH 89, a ∼1.8million-year-old partial hominid clavicle from Olduvai (Oldupai) Gorge, Tanzania. We compare the morphology and clavicular curvature of OH 89 to modern humans, extant apes, and a sample of other hominid fossil clavicles.

**Materials and Methods:** Comparative samples include 25 modern human clavicles, 30 *Gorilla*, 31 *Pan*, 7 *Papio*, and five hominid clavicles. Length regression on midshaft size using the extant comparative samples is used to estimate the total length of OH 89. A set of 9 linear measurements are taken from each individual. We also describe a new methodology for measuring clavicular curvature using measurements of sternal and acromial curvature, from which an overall curvature measurement is calculated. A principal component analysis (PCA) and a t-distributed stochastic neighbor embedding (tSNE) analysis are used to compare the morphology of OH 89 with the extant and fossil comparative samples.

**Results:** Our new method of measuring clavicular curvature successfully separates the different genera of the extant clavicles. The length estimate and sternal and acromial curve measurements for OH 89 falls within the larger male humans. The PCA shows OH 89 and most of the fossil hominids falling between the modern human and *Pan* groups, while the t-SNE suggests that OH 89, KSD-VP-1/1, KNM-ER 1808, and OH 48 are more similar to each other than to any of the other groups. This analysis also plots KNM-WT 15000 with the modern humans and Krapina 158 with the *Pan* individuals.

**Discussion:** The OH 89 clavicle derives from an individual of unknown hominid species with a shoulder breadth similar to that of a large human male. The curvature of OH 89 is relatively human-like relative to its length. Our new methodology for measuring clavicular curvature, combined with the utilization of t-SNE analyses and comparison of t-SNE results to PCA results, provides greater separation of genera than previously used methods, and wider use of t-SNE may be useful in paleoanthropological work.

## 1.0 Introduction

Until recently, paleoanthropologists did not dedicate much attention to the evolution of the hominid clavicle because its morphology is highly variable, and clavicles are relatively poorly preserved in the fossil record compared to other long bones. There are only a handful of decently preserved clavicles from Miocene apes, including a nearly complete *Nacholapithecus* clavicle from Kenya (Senut et al., 2004). The oldest hominid clavicle known to date is a fragment from *Ardipithecus* from the Afar region of Ethiopia (5.8-5.2Ma; Lovejoy, Suwa, Simpson, Matternes, & White, 2009). The Afar of Ethiopia has yielded a half dozen clavicular remains of *Australopithecus afarensis*, including a left clavicle associated with the partial skeleton KSD-VP-1/1 (3.6 Ma; Haile-Selassie et al., 2010; Melillo, 2016), both clavicles of a juvenile partial skeleton DIK-1-1 (Alemseged et al., 2006), and a handful of fragmentary clavicles from Hadar dating to 3.0-3.2Ma (Drapeau, Ward, Kimbel, Johanson, & Rak, 2005; Lovejoy, Johanson, & Coppens, 1982). *Australopithecus* clavicles are also known from South Africa. Sterkfontein has yielded partial clavicles StW 431g, StW 582, and StW 606, as well as the StW 573 specimen that preserves both clavicles of an *Australopithecus* individual dated to ca. 3.67 Ma (Carlson et al., 2021; Green, 2020). Two specimens of *Australopithecus sediba* (MH1 and MH2) from the site of Malapa also preserve incomplete clavicular fragments (1.95 Ma; Dirks et al., 2010). Although there are in total just over a dozen known clavicles of representing *Ardipithecus* and *Australopithecus*, these are mostly fragmentary and consequently, difficult to study.

As for the genus *Homo*, as we approach *H. sapiens* in time, the number of fossil clavicles increases. For example, KNM-WT-15000 from northern Kenya preserves nearly complete right and left clavicles of a juvenile *Homo erectus* (∼1.6 Ma; Walker, Leakey, & Leakey, 1993). KNM-WT 15000 has a notably short clavicle for its overall body size, though not outside of the range of modern human variation (Roach & Richmond, 2015). There are multiple partial clavicles attributed to *Homo naledi*, including U.W. 101-258, U.W. 101-1229, and U.W. 101-1347 (Feuerriegel et al., 2017). The majority of fossil clavicles are found in Europe, from Krapina, La Ferrassie, El Sidron, Gran Dolina, Sima de los Huesos, Kebara, Régardou, La Chapelle aux-Saints (Carretero, Arsuaga, & Lorenzo, 1997; Ferna et al., 1999; García-González, Rodríguez, Salazar-Fernández, Arsuaga, & Carretero, 2023; Gomez-Olivencia et al., 2018; Heim, 1982; RadovČić & Wolpoff, 1988; Rosas et al., 2016; Trinkaus, 2011; Voisin, 2006). Sima de las Palomas has also yielded a nearly complete Neanderthal clavicle (Walker et al., 2011). There are also roughly two dozen clavicles from Taforalt (Morocco), one from Qafzeh, and one from Shanidar (Trinkaus, 1982; Vandermeersch, 1981; Voisin 2006). There are multiple clavicles from Dmanisi, including the nearly complete D2724 (Lordkipanidze et al., 2007). For the earliest representatives of *Homo sapiens*, there is the Omo 1 partial skeleton that preserves a complete left clavicle from ∼130 ka (Day, 1969; Pearson, Royer, Grine, & Fleagle, 2008; Voisin, 2008).

The discovery of the clavicle from partial skeleton of *Australopithecus afarensis* KSD-VP-1/1 brought renewed attention to the clavicle and interpreting its role in the evolution of the hominid shoulder girdle (e.g., Melillo, 2016; Melillo et al., 2019). Applying new methodological approaches, Melillo (2016) concluded that all *Australopithecus* clavicular remains are indistinguishable from each other and that the clavicle of *A. afarensis* has morphological affinities to both chimpanzees and humans.

Although clavicles themselves do not provide as clear a taxonomic or functional signal as other aspects of the skeleton, the relatively conserved nature of the primate shoulder girdle does provide information about the evolution of primate locomotion in a broader sense. For example, from a much wider taxonomic perspective, the bones that make up the stylopod, zeugopod, and autopod of tetrapod forelimbs are fairly conserved while the skeletal structure of the girdle that attaches those forelimbs to the body varies widely. Bony fish have a scapula and a cleithrum, though its homology to the mammalian clavicle is debated (Shubin et al., 2015). Many amphibians have clavicles that are widely recognized as homologous to the mammalian condition, as well as a fused scapulocoricoid (Jenkins & Goslow, 1983). Reptiles, including birds, have scapulae that articulate with bony furculae that are fused at the inferior end (Nesbitt, Turner, Spaulding, Conrad, & Norell,2009). Many mammals possess clavicles to some degree, although they are often vestigial and essentially nonfunctional. The presence of a clavicle of some form across marsupials and placental mammals and other tetrapods suggests that the lack of a clavicle is a derived trait.

Because the clavicle acts as a strut to distribute forces from the arm to the axial skeleton, its presence and morphology reflect the biomechanical forces that act on the forelimb during different kinds of locomotion. A pair of clavicles in the shoulder girdle enables the upper limbs to move outside the parasagittal plane (Jenkins, 1974), in contrast to bovids and other cursorial animals who lack clavicles and are unable to move their forelimbs perpendicular to their spine. Within primates, brachiators, such as orangutans and gibbons, have absolutely and relatively long, slender clavicles that are laterally twisted along the long axis. This morphology is an adaptation to these primates’ superiorly oriented scapulae and tensile forces that act on the clavicles as the animals hang from their forelimbs. Arboreal quadrupedal primates, such as cercopithecins and colobines, have short, j-shaped clavicles and a deep fossa on the inferior side of the acromial end. As for fossils, Melillo (2016) interpreted *Australopithecus afarensis* clavicular morphology as evidence of increased manipulatory function of the upper limb.

The clavicle of *Homo sapiens* is notoriously variable and may well have the most developmental plasticity of any bone in the human skeleton. It is the first to begin ossification at around 9 weeks in utero and is one of the last bones to fully fuse between 20 and 30 years in age (Black & Scheuer, 1996). Developmentally, the clavicle is unique in having two primary ossification centers with different developmental origins. The medial end develops endochondrally from embryonic mesoderm while the lateral end develops intramembranously from embryonic neural crest (Matsuoka et al., 2005; Ogata & Uthoff, 1990). The extensive duration of ontogenetic development acting on these two centers of ossification means that the accumulation of biomechanical forces acting upon the shoulder girdle from before birth well into adulthood can influence adult clavicle morphology. Given this long ontogeny of the clavicle, the evolution of its morphology may well provide us with a window into a long period of the genetic and epigenetic forces experienced by the forelimbs of hominid individuals over geological time in addition to the broader insight to locomotory behavior.

In 2005, the lateral end of a hominid clavicle was recovered by AVS and other members of the Conservation Olduvai Project from Olduvai (Oldupai) Gorge, Tanzania. Following the historical practice of sequentially numbering hominid fossils from Olduvai Gorge, this clavicle is designated OH 89 (Figure 1). This is the second hominid clavicle discovered at Olduvai Gorge. The first, OH 48 was originally associated with the ∼1.83 million-year-old OH 8, then later given its own number as its association with OH 8 was unclear (Day & Scheuer, 1989). OH 89 (Figure 1) preserves the lateral half of a right hominid clavicle.

**Figure 1.**
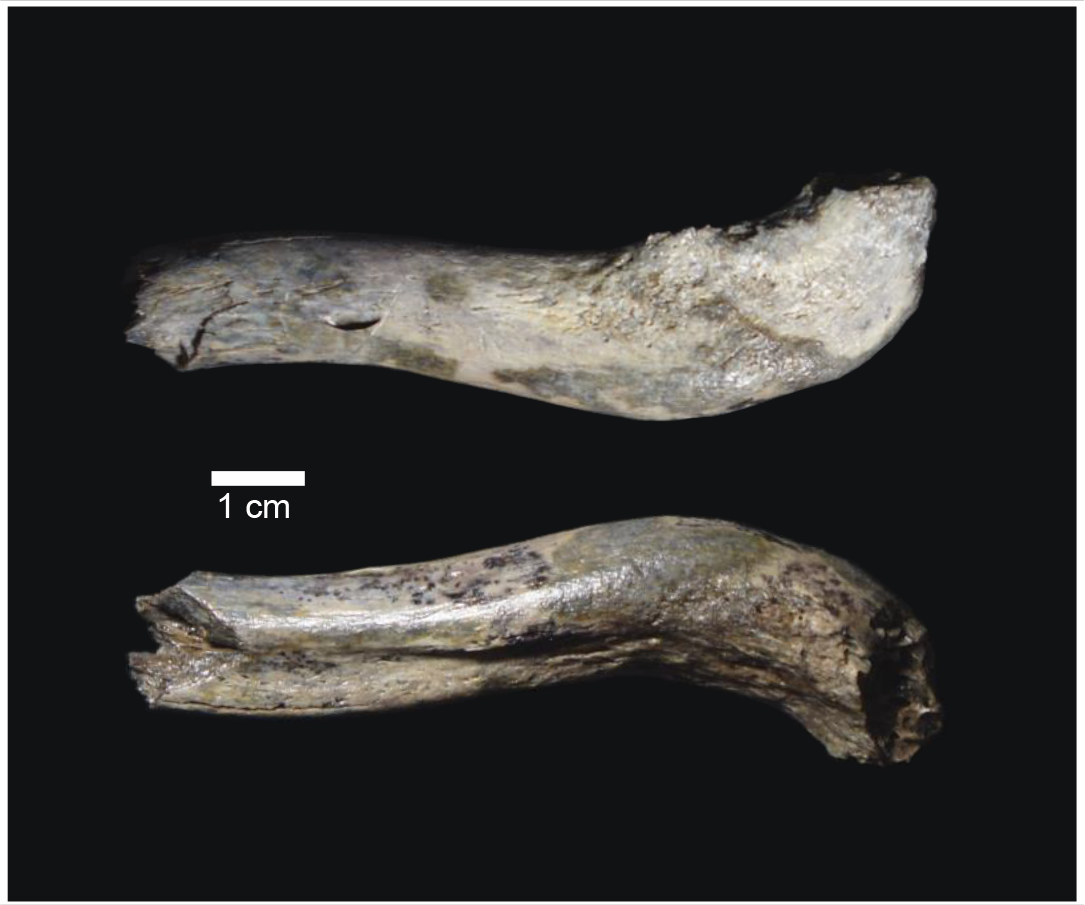
OH 89 in superior view (top) and inferior view (bottom).

OH 89 was found on the surface near the FLK North locality, eroding from upper Bed 1, just below tuff 1F (Figure 2). Tuff 1F has most recently been ^40^Ar/^39^Ar dated to 1.803 million years ago (Ma; Deino, 2012; Deino et al., 2021). Therefore, the minimum age of OH 89 is determined to be ∼1.8 Ma. Given this geological age, OH 89 represents an important point of comparison between the earlier eastern African *A. afarensis* clavicles at 3.5 Ma, the approximately 1.95 Ma clavicular specimens from *Australopithecus sediba*, the 1.6 Ma clavicle of *Homo erectus* (KNM-WT-15000), and the clavicles of the more recent representatives of *Homo* from the Middle East and Eurasia. This time period is especially important given the morphological changes that occurred between *Australopithecus* and the appearance of *Homo*. The OH 89 clavicle, although isolated and fragmentary, provides an important data point in understanding the evolution of the hominid shoulder. OH 89 is now housed at the Arusha National Natural History Museum in Arusha, Tanzania.

**Figure 2.**
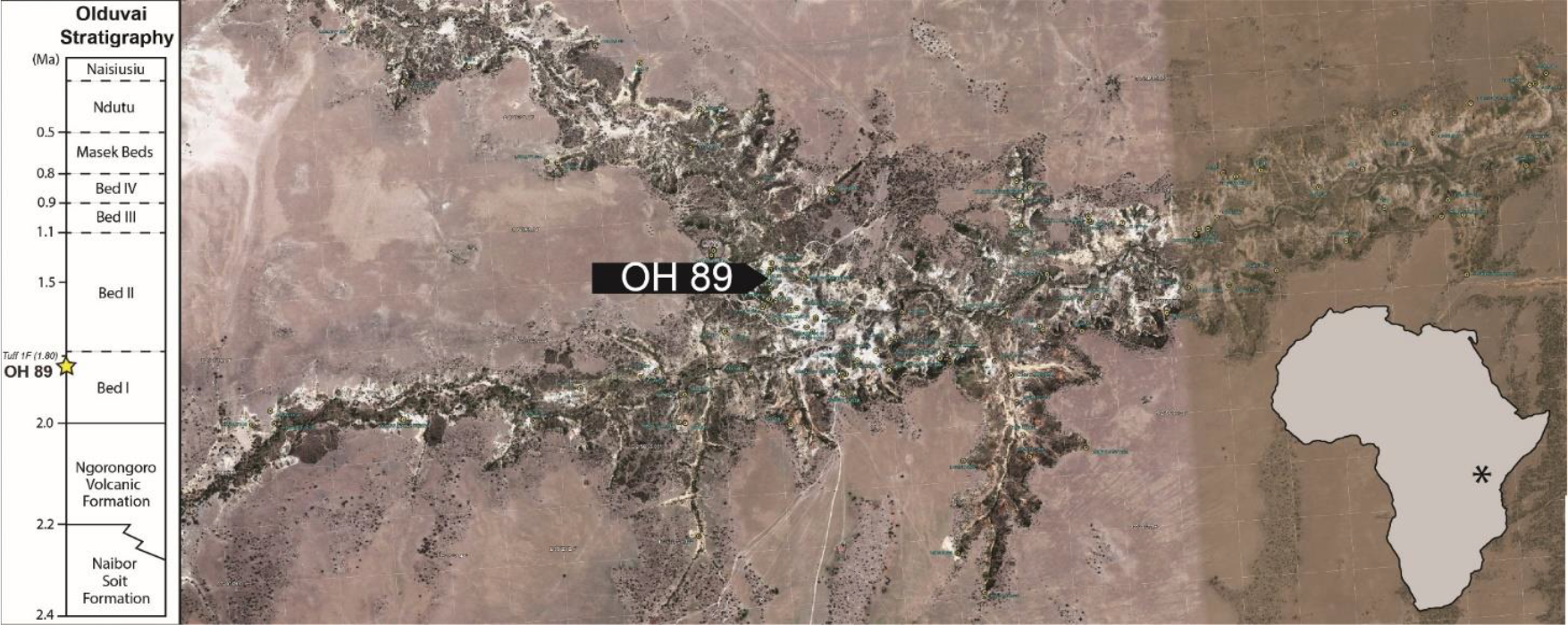
Map showing the location of OH 89 at Olduvai Gorge and the stratigraphic position of the fossil (left), with a map of Africa showing the geographic location of Olduvai Gorge (right). The stratigraphy is based on Hay (1976), Deino (2012), Deino et al. (2021), Deiz-Martín et al. (2015), McHenry & Stanistreet (2018), Njau et al. (2021), and Stanistreet et al. (2020).

## 2.0 Preservation and Anatomical Description of OH 89

OH 89 (Figure 1) has a maximum length of 87.9 mm and a maximum anteroposterior width of 20.4 mm. The acromial articular surface is missing, and the clavicle is broken around midshaft. There is a distinct subclavian groove, and the clavicle is roughly crescent shaped in cross section near the break. The rugosity for the deltoid is distinct and relatively more robust than that of the trapezius. The conoid tubercle appears as more of a rugosity than a tubercle.

## 3.0 Materials

We compared OH 89 to a sample of extant human and non-human primate and fossil hominid clavicles. Our extant sample consists of adult individuals from *Gorilla gorilla* (n=30), *Pan troglodytes* (n=31), *Homo sapiens* (n=25), and *Papio hamadryas* (n=7). We only included specimens that had no evidence of trauma or pathology. All extant non-human individuals come from wild-shot animals that are part of the Hamann-Todd Osteological Collection housed at the Cleveland Museum of Natural History (CMNH).

All 25 modern human individuals are housed in the Hamann-Todd Collection at CMNH. These skeletal remains are from people who died in the Cleveland area in the late 19^th^ and early 20^th^ centuries and whose bodies were unclaimed from hospitals and morgues (Cobb, 1933). These people therefore disproportionately represent low-income groups and, given the standards of medical ethics at the time, most likely did not agree to donate their remains to science.

All of these skeletal collections present complicated ethical questions (e.g., Walker, 2000; de la Cova, 2019; Squires and García-Mancuso, 2021). As our aim for this research is to better understand human and non-human primate biology from the broad perspective of geological time, we intentionally do not consider the ethnic or personal identification information of anyone included in this study. For the people whose clavicles were measured as part of in our study, we deeply appreciate their inclusion as representatives of *Homo sapiens*. For the non-human primates included in this study, we hope that using their remains to better understand primate biology at least derives some good from the sacrifice of wild animals.

Fossil clavicles included in this study are shown in Table 1. All measurements from fossils were taken on research quality casts of the original fossils, housed in the Human Evolution Research Center at UC Berkeley and the Cleveland Museum of Natural History.

**Table 1.** Fossil clavicles included in analyses.

| Specimen Number | Side | Species | Date | References |
| --- | --- | --- | --- | --- |
| KNM-WT 15000 | R | *Homo erectus* | 1.53 Ma | Walker & Leakey, 1993 |
| KNM-ER 1808 | R | *Homo erectus* | ~1.5 Ma | Walker, Zimmerman & Leakey, 1982 |
| OH 48 | L | Unidentified | 1.84 Ma | Napier, 1965 |
| Krapina 158 | L | Neanderthal | ~130 Ka | Rink et al., 1995 |
| KSD-VP-1/1 | L | *Australopithecus afarensis* | 3.6 Ma | Haile-Selassie et al., 2015 |

## 4.0 Methods

### 4.1 Linear Measurements

A series of 9 linear measurements were taken using Mitutoyo digital calipers: Maximum width of acromial end, maximum anteroposterior shaft width, maximum anteroposterior dimension, length of acromial rugosity, superoinferior width at midshaft, anteroposterior width at midshaft maximum superoinferior height of acromial region, anteroposterior width at conoid tubercle, and superoinferior width at conoid tubercle. Additionally, maximum length measurements were taken using an osteometric board. All linear measurements were taken directly on original specimens (for extant samples) or research quality casts (for fossil samples).

### 4.2 Length Estimate

Bilateral asymmetry of anthropoid clavicles is well documented and results in right and left clavicles of the same individual exhibiting different lengths (Parsons, 1916; Schultz, 1937; Ray, 1959; Mays, Steele, & Ford, 1999; Voisin, 2001; Auerbach and Raxter, 2008; Abdel Fatah et al., 2012). We calculated an estimated length for OH 89 based off of the right clavicle using an ordinary least squares regression on midshaft size (√minimum midshaft width x perpendicular measurement), following Melillo (2016). These estimates use *Gorilla, Pan* and modern human individuals as a reference sample.

### 4.3 Curvature Measurement

Many researchers have quantified clavicular curvature through a range of different approaches, all of which have pros and cons (Olivier, 1951; Squyers & DeLeon, 2006; Voisin, 2006; Melillo, 2016; Barros, 2014). Linear measurements can quantify magnitude of curvature, but such analyses entirely lose the taxonomic shape profiles. Linear analyses are also unable to distinguish between a long clavicle with shallow curves and a short clavicle with deep curves. Alternatively, 3D geometric morphometrics can be especially difficult to apply reliably to the clavicle because there are virtually no landmarks that are consistent across taxa, and those that are present are highly variable even within one species. Semi-landmarks may be utilized, but the reproducibility and accuracy of landmark placement is low and therefore less than ideal. 2D geometric morphometrics has most accurately captured clavicular shape and curvature thus far, but still has not been able to quantified the morphological differences as well as can be done by eye.

The methodology used here to assess clavicular curvature is partially modeled after that of Melillo (2016) but uses a simpler geometric approach instead of 2D geometric morphometric analyses. Given the preservation of OH 89, we only assess curvature from superior view. All specimens were photographed in a standard superior position with the acromial end horizontal to the ground. A curvilinear mid-line was drawn for each specimen that represents the sinusoidal shape of the bone. A composite is presented in Figure 3 to give the reader a sense of the range of variation in sinusoidal curvature in each of the taxa included in this study.

**Figure 3.**
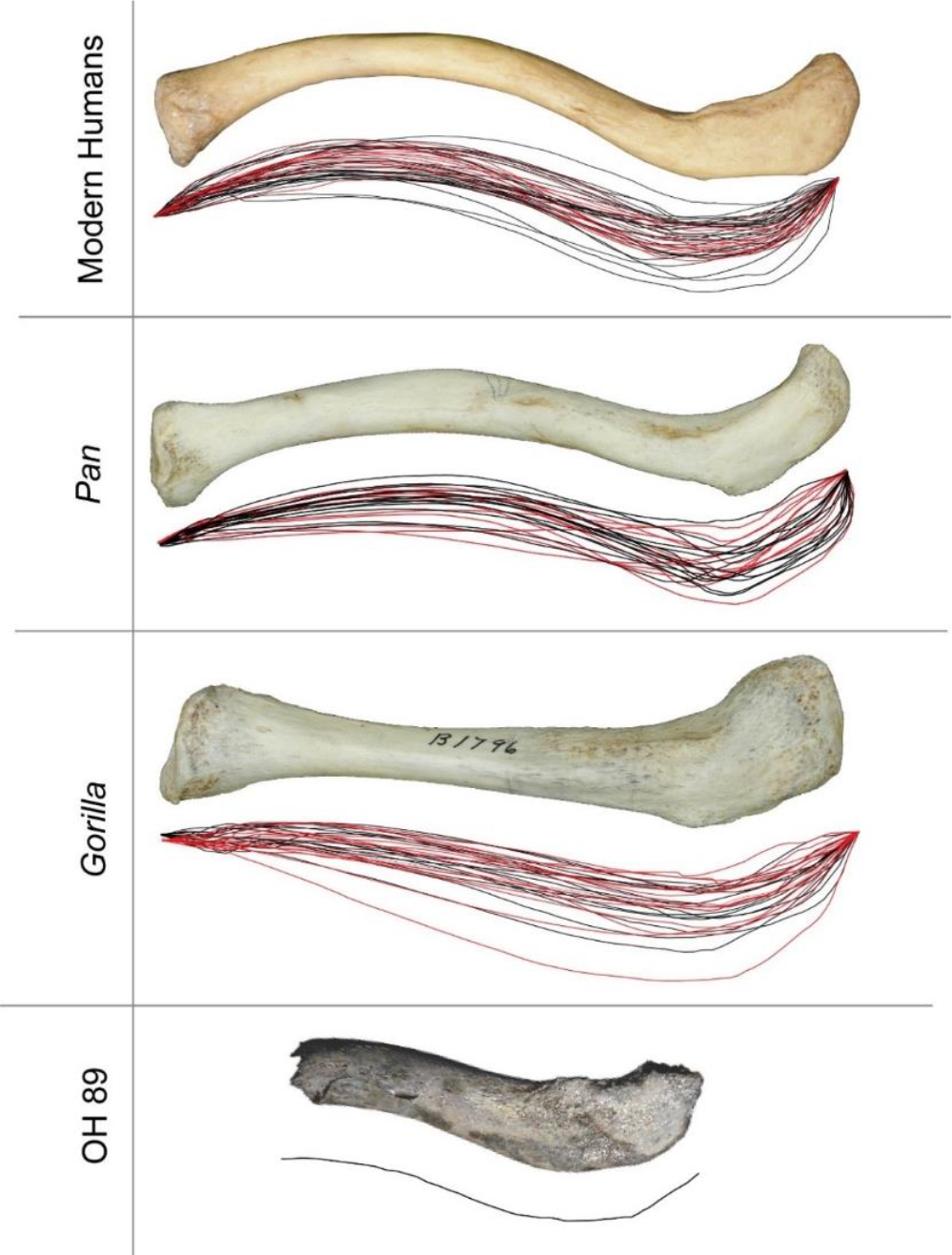
Visualization of variation in clavicular curvature in comparative samples. Each midline curve was scaled to the same size and overlaid with others of the same taxon to create a “curvature cloud”. This visualization allows us to see the differences in overall shape of clavicular curvature between taxa as well as the variation within a group. All photos are scaled to the same size to better facilitate comprehension of shape without relation to size. Red lines indicate a female individual and black lines indicate a male individual.

Measurements of curves were taken in Adobe Illustrator using the following steps (illustrated in Figure 4):

1. Midline curves were drawn on standardized photos of clavicles following the methodology of Melillo (2016).
2. The two endpoints of each midline curve were fitted to a horizontal line. The sternal and acromial endpoints are defined as the terminal midline point on each end of the clavicle.
3. Two circles were overlain on each clavicle using the “ellipse” tool in Adobe Illustrator, one on the sternal and one on the acromial curve, such that the circle touches both endpoints of each curve and the maximum height of the curve.
4. A measurement of the radius of the sternal and acromial circles was taken. Midpoints for each circle were geometrically determined in Adobe Illustrator, and radius of each circle were automatically calculated by Adobe Illustrator in pixels, then converted to millimeters.

**Figure 4.**
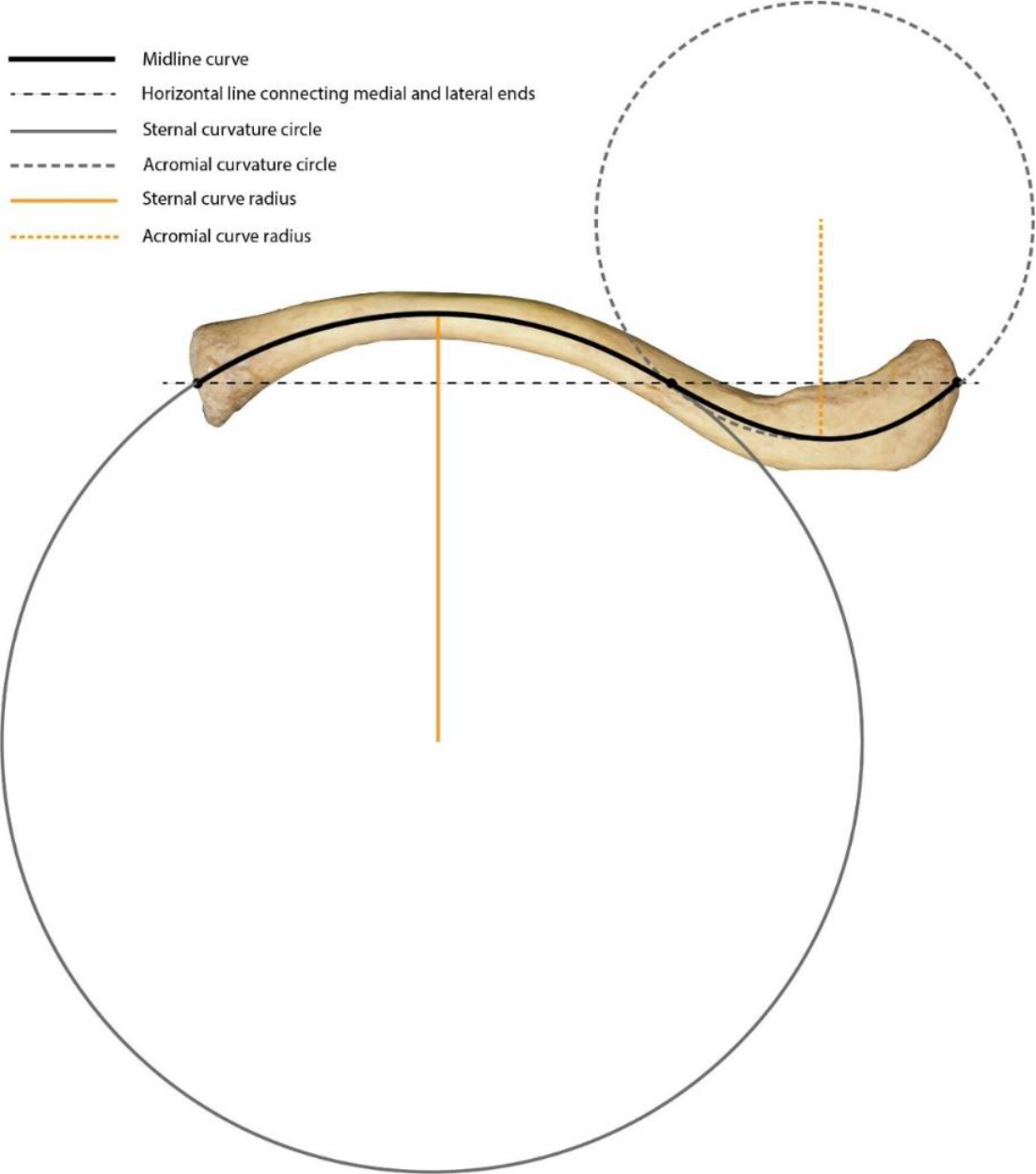
An illustration of the methodology used to study curvature. See text for details.

These two curves (radius of sternal curve and radius of acromial curve) were then used to calculate a ratio of radii. Not only does this method capture both the length and depth of each curve, but it also allows for easier visualization of phylogenetic differences in morphology (see Results). Data are available at https://doi.org/10.6084/m9.figshare.22050185.v1

### 4.4 Analyses

We ran a series of standard univariate and bivariate analyses that include univariate descriptive statistics, a bivariate plot, and a principal component analysis (PCA). Univariate descriptive statistics were estimated using the *tapply* function of the *IRanges* package (Lawrence et al., 2013). Plots were produced using *ggplot2* (v3.3.0, Wickham, 2016) in the R statistical environment v3.6.1 (R Core Team, 2019) and edited in Adobe Illustrator CC 2019 (Adobe, Inc.). PCA includes all 11 measurements taken (the 9 linear measurements listed in section 4.1 and the two curve radius measurements described in section 4.3) and was performed using the *plotly* package (Sievert, 2020).

We also include a relatively new method of analysis for morphological studies – a t-distributed stochastic neighbor embedding analysis (t-SNE). This analysis was run using the *rtsne* package in R with a perplexity of 35 and 200 steps. t-SNE is a technique of visualizing high-dimensional data in two dimensions, somewhat similar to a PCA but with the capability of collapsing multi-dimensional data into a two-dimensional visualization (Krijthe, 2015; van der Maaten & Hinton, 2008). R code used in analyses can be found at https://doi.org/10.6084/m9.figshare.22054280.v1.

The goal of including this analysis was to determine if its use provides any additional information about fossil morphology when compared to a PCA. While the two analyses provide a similar way of visualizing data, t-SNE analyses collapse all of the data into two dimensions, rather than picking out two individual variables from the data and discarding the rest of the variables.

## 5.0 Results

Univariate descriptive statistics are presented in Table 2. To visualize the variation within and between taxa, maximum length is plotted against midshaft size for all extant and fossil data in Figure 5. Our length regression on midshaft size analysis estimates that OH 89 had a maximum length of 155.6 ± 2.4 mm. The fitted regression model was: Maximum length = 38.1713 + 9.3131 * Midshaft size (R^2^ = 0.73, p < 0.001). Based on our regression on midshaft size, we estimate that the total length of OH 89 falls toward the upper end of modern human male variation in both maximum length and midshaft size. Compared to the other fossil clavicles included in our study, OH 89 is the largest in all dimensions.

**Table 2.** Descriptive statistics of fossils and extant comparative samples. Ranges are displayed as minimum value to maximum value. SI = Superoinferior and AP = Anteroposterior. All values are in mm except the ratio of radii. Ratio of radii is defined as the radius of the sternal curve divided by the radius of acromial curve.

|  |  | Maximum Length | Maximum width of acromial end | Minimum AP Shaft Width | Maximum AP Dimension | Superoinferior Width at Midshaft | Maximum Superoinferior Height of Acromial Region | Anteroposterior Width at Conoid Tubercle | Superoinferior Width at Conoid Tubercle | Anteroposterior Width at Midshaft | Radius of Internal Curve | Radius of External Curve |
| --- | --- | --- | --- | --- | --- | --- | --- | --- | --- | --- | --- | --- |
| *G. gorilla* (n = 30) | Mean | 153.2 | 34 | 12.7 | 66.6 | 11.1 | 11.2 | 21.3 | 11.3 | 13.7 | 41.9 | 145.02 |
|  | Range | 121.4 - 184.7 | 22.5 - 46.5 | 8.9 - 18.1 | 53.8 - 79.0 | 8.8 - 15.4 | 7.3 - 17.8 | 15.8 - 27.6 | 8.4 - 15.1 | 9.4 - 17.7 | 0 - 173.3 | 81.5 - 283.6 |
| *P. troglodytes* (n = 31) | Mean | 127.5 | 29.3 | 10.5 | 54.8 | 8.3 | 11.8 | 17.3 | 8.6 | 12.2 | 120.5 | 39.7 |
|  | Range | 113.1 - 146.6 | 23.0 - 34.9 | 9.2 - 11.7 | 28.2 - 60.9 | 6.4 - 9.8 | 9.2 - 14.3 | 12.7 - 22.1 | 6.9 - 10.7 | 9.5 - 14.5 | 71.5 - 283.8 | 26.2 - 66.3 |
| *H. sapiens* (n = 25) | Mean | 144.2 | 31.5 | 11.6 | 62.5 | 9.8 | 11.4 | 17.7 | 10.9 | 12.1 | 91.6 | 49.1 |
|  | Range | 127.4 - 158.8 | 26.2 - 37.0 | 8.7 - 14.0 | 54.8 - 68.6 | 7.8 - 13.1 | 7.9 - 14.5 | 15.1 - 21.9 | 8.3 - 15.2 | 9.0 - 14.8 | 69.4 - 135.8 | 29.9 - 74.0 |
| *P. hamadryas* (n = 7) | Mean | 73.24 | 20.1 | 6.3 | 33.2 | 5.7 | 6.9 | 8.8 | 5.2 | 6.7 | 2.6 | 47.5 |
|  | Range |  | 13.4 - 26.3 | 4.6 - 8.8 | 24.6 - 43.9 | 4.6 - 7.4 | 5.2 - 8.8 | 5.5 - 14.2 | 4.3 - 6.7 | 4.5 - 8.6 | 0 - 15.6 | 30.7 - 60.3 |
| OH 48 | Mean | 148 | 16.2 | 12.3 | 15.3 | 10.3 | 11.3 | 20.7 | 11.8 | 15.2 | 108.2 | 36.2 |
|  | Range | - | - | - | - | - | - | - | - | - | - | - |
| OH 89 | Mean | 155.7 | 20.04 | 14.8 | 19.8 | 10.5 | 11.1 | 15.8 | 9.5 | 9.4 | 54.6 | 38.9 |
|  | Range | 153.3 - 158.0* | - | - | - | - | - | - | - | - | - | - |
| KNM-ER 1808 | Mean | 142.8 | 14.7 | 10.6 | 12.3 | 9.3 | 9 | 15.8 | 8 | 12.2 | 90.2 | 35.4 |
|  | Range | - | - | - | - | - | - | - | - | - | - | - |
| KNM-WT 15000 | Mean | 130.5 | 22.4 | 10.6 | 54.5 | 8.9 | 9 | 14.1 | 8.8 | 10 | 87.2 | 44.3 |
|  | Range | - | - | - | - | - | - | - | - | - | - | - |
| Krapina 158 | Mean | 126.7 | 28.5 | 15.6 | 17.3 | 8.8 | 9.8 | 18.4 | 9 | 11.3 | 151.1 | 51.5 |
|  | Range | - | - | - | - | - | - | - | - | - | - | - |
| KSD-VP-1/1 | Mean | 157.2 | 30.6 | 15.3 | 24.6 | 9.6 | 10 | 20.3 | 10.4 | 16.4 | 62.2 | 43 |
|  | Range | - | - | - | - | - | - | - | - | - | - | - |

**Figure 5.**
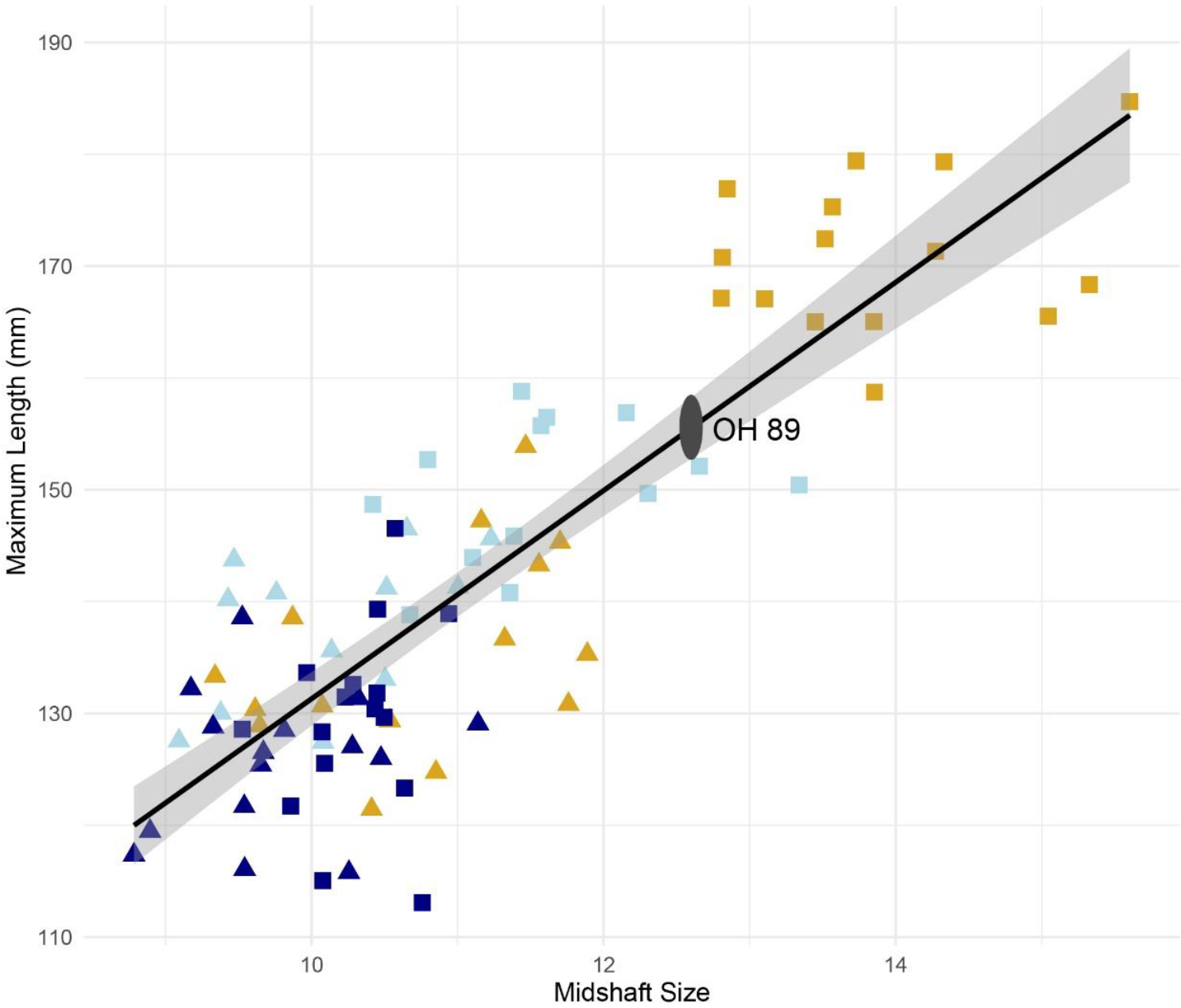
A length regression on midshaft size based on the right clavicles of modern humans, *Gorilla*, and *Pan* individuals. *Pan* individuals are in dark blue, modern humans are in light blue, and *Gorilla* are in gold. Triangles indicate female and squares are male. OH 89’s length estimate based on midshaft size is indicated by the grey oval, with a mean of 155.7 mm.

Our quantification of clavicular curvature using circles captures the variation observed in bivariate plots (Figure 6), and PCA and t-SNE (Figure 7). For bivariate plots, both the ratio of radii and the radius of the acromial curve are used. Considering the fragmentary preservation of OH 89, we provide our estimate based on reconstructed length of the sternal curve, but error is high due to preservation, so the acromial curve is presented as well. While there is some overlap between humans and *Pan*, and humans and *Gorilla*, the three groups are mostly distinct. The PCA (Figure 7, left) appears to capture size and curvature in a similar manner to the bivariate plot (Figure 6). This quantitatively demonstrates that size and curvature profiles are the two most distinguishing characters in clavicles, according with the pattern discernible by eye.

**Figure 6.**
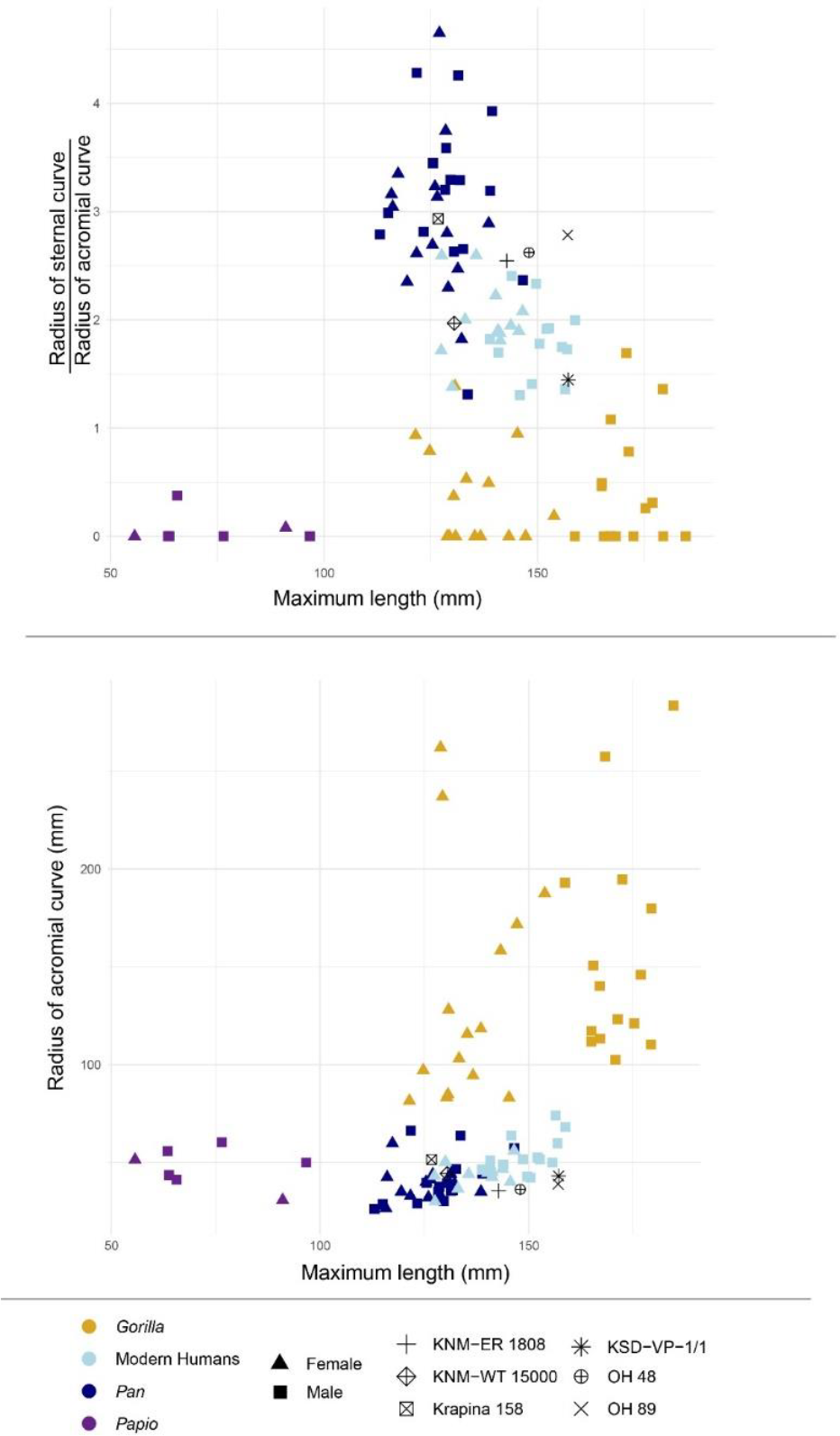
Bivariate plots showing the ratio of radii vs. maximum length (top) and the radius of the acromial curve vs. maximum length (bottom).

**Figure 7.**
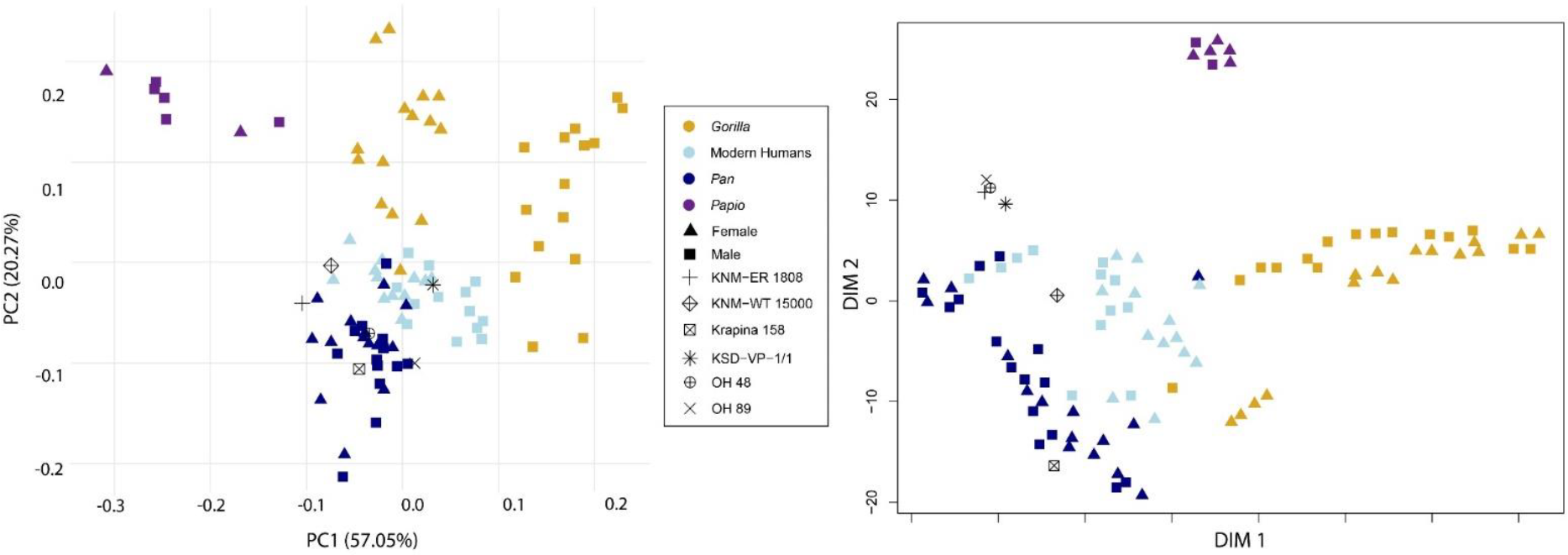
Left: Principal components analysis (PCA) showing fossil and extant clavicles. Right: t-distributed stochastic neighbor embedding (t-SNE) showing fossil and extant clavicles.

Turning to the fossil data, we note that the fragmentary preservation of OH 89 requires that we interpret results with caution. With that said, the PCA shows minor overlap between chimpanzee and human individuals, with the hominid fossils falling within the two groups (Figure 7, left panel). OH 89 falls on the edge of the range of human variation in acromial curvature, and right on the edge between human and chimpanzee variation in the ratio of radii (Figure 6). Krapina 158 differs most apparently from *Homo sapiens* in comparison to the other fossils. The t-SNE results demonstrate a close affinity between humans and chimpanzees (Figure 7, right panel). In t-SNE analyses, the distance between clusters is not necessarily as informative as the clusters themselves. Thus, the plotting of several fossils on the top left does not necessarily indicate that those individuals are more similar to chimpanzees than humans, but does suggest that those individuals are more similar to each other than to any of the other groups. In this case, Figure 7, right panel shows that KNM-WT 15000 is most similar to humans, Krapina 158 to *Pan*, and that KNM-ER 1808, KSD-VP-1/1, OH 48, and OH 89 are most similar to each other.

## 6.0 Discussion

Analyses of complete hominid clavicles have provided justification for claims about shoulder height, throwing ability, manual dexterity, and climbing ability (e.g., Larson, 2007; Roach & Richmond, 2015). The bilateral asymmetry of clavicles offers particular insight to behaviors, past and present (e.g., Abdel Fatah et al., 2012; Auerbach & Raxter, 2008; Trinkaus, Churchill, & Ruff, 1994). Given this relatively high degree of epigenetic influences on clavicular morphology compared to other regions of the skeleton, a fragmentary clavicle is a frustrating fossil to discover in isolation. Such a fossil is difficult to interpret taxonomically, and out of skeletal context, difficult to utilize in paleobehavioral studies. However, there is still information to be learned about human evolution from these fossils.

Our analysis of the fragmentary clavicle OH 89 provides clear evidence that a hominid died in the region of now-Olduvai Gorge 1.8 million years ago. This individual’s clavicle was at the larger end of the variation we observe in *Homo sapiens* and is the largest of the hominid clavicles included in our analysis. While clavicle length is not a good proxy for body size, it is a good predictor of the breadth of the individual’s shoulders (Melillo et al., 2019). Our results indicate that OH 89 derives from an individual with a shoulder breadth similar to a large male today. With larger human comparative sample sizes, OH 89 would likely fall within the range of large humans.

One of the most defining morphological characteristics of the clavicle is its curvature. While this curvature is distinctive to the human eye (making the differentiation of hominoid clavicles fairly straight-forward qualtitatively), quantifying these shape differences has been difficult. Multiple studies have attempted at quantifying clavicle curvature (Olivier, 1951; Voisin, 2006; Daruwalla, Courtis, Fitzpatrick, Fitzpatrick, & Millett, 2010; Abdel Fatah et al., 2012 ; Bernat et al., 2014; Squyers & DeLeon, 2015; Melillo, 2016; Barros, 2014). Here, we introduced a new method designed to be more easily applied to fragmentary fossils. Our method uses two circles to quantitatively capture the relative sinusoidal curvature of the clavicle. We find that the estimated sinusoidal curvature of OH 89’s clavicle, as well as that of OH 48, is human-like relative to its length.

But OH 89 and OH 48 are clearly not identical to humans. Our PCA shows OH 89 and OH 48 at the intersection between humans and chimpanzees, along with KNM-WT 15000. A reasonable interpretation is that these three fossils are all part of the *Homo erectus* morphology, that chimpanzee clavicular morphology is ancestral and human morphology is derived. However, this interpretation is unsupported when we consider the anatomy of KSD-VP-1/1 which falls in the center of human variation (similar to Melillo’s observation, 2016), and that the Neanderthal morphology of Krapina 158 is like that of chimpanzees. Researchers have noted the unique morphology of Neanderthal clavicles previously, so this aspect of our results is not entirely surprising (Mersey, Brudvick, Black, & Defleur, 2013; Voisin, 2006; Trinkaus, Holliday, & Auerbach, 2014; Rosas et al., 2016). However, the human-like morphology of the oldest clavicle included here (KSD-VP-1/1, 3.6 Ma), and the somewhat offset morphology of the 1.5 – 2 Ma clavicles suggest that evolution of the hominid clavicle has not been a constant movement towards a modern human morphology from a more chimpanzee morphology. This result echoes the conclusions reached by investigations of *Ardipithecus* anatomy (Lovejoy et al., 2009; Lovejoy and McCollum, 2010; White et al., 2015), showing that chimpanzees are distinctly derived compared to the chimpanzee-human last common ancestor (LCA) and therefore not an appropriate stand-in for the chimpanzee-human LCA.

Our t-SNE analysis, which relies on machine-learning (Figure 7), converts high-dimensional data into pairwise similarities, and thereby provides useful data visualization (Arora, Hu, & Kothari, 2018). The results of t-SNE analyses are designed to provide visual insight into the patterns formed by a data set, not provide quantitative results about the similarities and differences between groups. This type of analysis is usually employed in other disciplines with larger data sets, but it does provide insight into fossil morphology as illustrated here. We chose to utilize this analysis in an effort to determine if the conclusions of a t-SNE would be similar to a PCA, or if this analysis could provide further insight that the PCA does not. We found that most of the hominid clavicles cluster with each other, suggesting that they are more similar to each other than they are to any other clavicle included in the analysis. The two notable outliers to the hominid cluster are Krapina 158 and KNM-WT 15000. The Krapina clavicle is also somewhat of an outlier compared to the other hominids in its curvature profile, with a relatively larger sternal curve compared to the other fossils (Figure 6). Krapina 158 is also the shortest of the fossil clavicles, followed closely by KNM-WT 15000. The placement of KNM-WT 15000 closest to modern human in the t-SNE may be driven by the small size of this individual, given its sub-adulthood.

There are caveats to t-SNE, not the least of which is that it is usually used for big data sets with dozens to hundreds of variables, and therefore may be inappropriate for these much smaller datasets. However, it is fascinating that t-SNE returns an output that differentiates between our samples better than previously published analytical methods, and that these results compare favorably with what is visually obvious: human, chimpanzee, and gorilla clavicles look different from each other, KNM-WT 15000 falls most closely with modern humans, and that Krapina 158 is distinct from humans and other fossil hominids. These results support the conclusion by Voisin (2008) that OH 48 and KNM-WT 15000 have somewhat more pronounced sternal curves than acromial curves, though the results here place those two individuals on the edge of human variation.

In summary, our analyses demonstrate that OH 89 is, for the most part, on the edge of the range of variation of human clavicular variation despite its antiquity. Although researchers occasionally assign fragmentary, isolated postcrania to species, such a taxonomic identification is not possible for OH 89. Based on other fossil evidence, we know that around ∼1.8 Ma, there were at least two distinct species of hominids living around Olduvai Gorge (Blumenschine et al., 2012; Clarke, 2012; Domínguez-Rodrigo et al., 2015). While it is impossible to decisively say what species OH 89 and OH 48 belonged to, or if they are even conspecific, the discovery of OH 89 does reveal that hominid clavicles show notable similarities to modern humans at least as far back as 1.8 Ma. This finding suggests that the evolutionary and biomechanical forces acting on the arms and shoulders of 1.8-million-year-old hominids are not entirely dissimilar to those acting on modern humans today.

## 7.0 Conclusion

OH 89, a 1.8 Ma hominid clavicle, falls within the range of modern humans in absolute size and clavicular curvature. This finding indicates that there has been little morphological change in the hominid clavicle in the last ∼2 million years. It also suggests that shoulder breadth (though not necessarily body size) may have been similar to modern humans as far back as 1.8 Ma.

## Acknowledgements

This project would not have been possible without the support and assistance of: Dr. H. Mshinda, Director General, Commission for Science and Technology (COSTECH); Mr. J. Paresso and Mr. J. Temba, Tanzanian Department of Antiquities; Ms. V. Ufunguo, Ngorongoro Conservation Area Authority; Dr. A. Mabulla and Mr. J. Kihiyo, current and past directors of the National Museum of Tanzania; Dr. P. Msemwa, Dr. A. Kweka, and Dr. A. Gidna, Museum and House of Culture, Dar es Salaam, Tanzania; Mrs. F. Mangalu, Director of the National Natural History Museum, Arusha; OVPP’s Olduvai field crew; Elias Bura, Theresa Grieco, Maneno Hussein, Odee Lyimo, Dan Mainoya, Justin Steven, Frank Mataro, Harry Mlaki, Leken Olle Moita, Mateo Oltusi, Singira Oltusi, Whitney Reiner, Kerio Zebedayo, Nairoshi Zebedayo, Richard Bakari, and Ikoyo Singoi (aka Bruce Lee). Many thanks to the museums and staff who facilitate our access to the comparative skeletal collections, including L. Jellema from CMNH and N. Johnson from the Hearst Museum for approving access. Additionally, thank you to Yohannes Haile-Selassie for providing access to KSD-VP 1/1.

## Data Availability

Data used in this study are available at https://doi.org/10.6084/m9.figshare.22050185.v1. R code is available at https://doi.org/10.6084/m9.figshare.22054280.v1.

